# RNA-bound PGC-1α controls gene expression in liquid-like nuclear condensates

**DOI:** 10.1101/2020.09.23.310623

**Authors:** Joaquín Pérez-Schindler, Bastian Kohl, Konstantin Schneider-Heieck, Volkan Adak, Julien Delezie, Geraldine Maier, Thomas Sakoparnig, Elyzabeth Vargas-Fernández, Bettina Karrer-Cardel, Danilo Ritz, Alexander Schmidt, Maria Hondele, Sebastian Hiller, Christoph Handschin

## Abstract

The peroxisome-proliferator-activated receptor-γ coactivator-1α (PGC-1α) integrates environmental cues by controlling complex transcriptional networks in various metabolically active tissues. However, it is unclear how a transcriptional coregulator coordinates dynamic biological programs in response to multifaceted stimuli such as endurance training or fasting. Here, we discovered a central function of the poorly understood C-terminal domain (CTD) of PGC-1α to bind RNAs and assemble multi-protein complexes. Surprisingly, in addition to controlling the coupling of transcription and processing of target genes, RNA binding is indispensable for the recruitment of PGC-1α to chromatin into liquid-like nuclear condensates, which compartmentalize and regulate active transcription. These results demonstrate a hitherto unsuspected molecular mechanism by which complexity in the regulation of large transcriptional networks by PGC-1α is achieved. These findings are not only essential for the basic understanding of transcriptional coregulator-driven control of biological programs, but will also help to devise new strategies to modulate these processes in pathological contexts in which PGC-1α function is dysregulated, such as type 2 diabetes, cardiovascular diseases or skeletal muscle wasting.

## Main

PGC-1α controls complex biological processes by regulating transcriptional networks sensitive to environmental cues such as fasting in liver, cold in brown adipose tissue and exercise in skeletal muscle^1-3^. Indeed, skeletal muscle-specific overexpression of PGC-1α in mice induces an endurance trained-like phenotype, characterised by robust mitochondrial biogenesis and enhanced exercise performance^1,4^. The molecular underpinnings that enable such a broad transcriptional control in a spatiotemporal manner however are unclear. PGC-1α lacks any discernible enzymatic activity and its function is thought to depend on the formation of a multi-protein complex containing a multitude of different transcription factors (TFs) and additional transcriptional regulators^5,6^. PGC-1α is highly conserved from mouse to human with 94% identity. The protein is intrinsically disordered for the large majority of its 797 amino acids and contains only a single folded domain, an RNA recognition motif (RRM) (Fig. 1a, Fig. 4c). Three sequence features are present in the intrinsically disordered part, an LXXLL motif, typically involved in nuclear receptor binding^7^, and two arginine–serine-rich (RS) regions. The segment of residues 564–797, comprising the two RS and the RRM, is generally termed the C-terminal domain (CTD). Deletion of the CTD of PGC-1α blunts its positive effects on gene expression, which was originally attributed to impaired RNA processing^8,9^. A direct coupling of transcription to RNA processing was underlined by PGC-1α interactions with splicing factors and components of the RNA polymerase 2 (Pol 2) complex via the CTD and a mini-gene splicing assay^8^. However, the impact of PGC-1α on RNA binding and processing of target genes and, hence, the functional relevance of the CTD remain unclear.

**Fig. 1:**
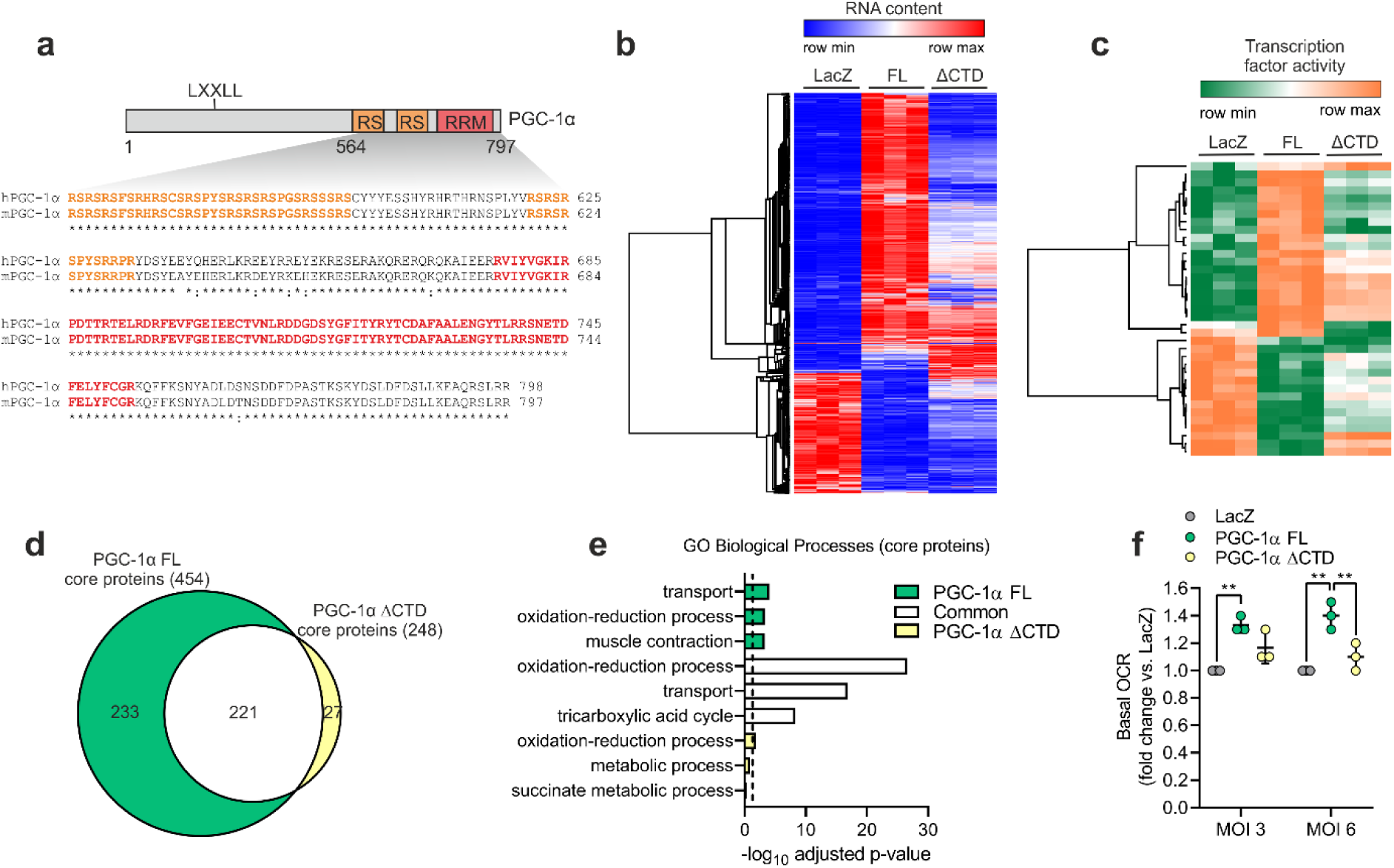
PGC-1α function is impaired by deletion of the CTD. **a**, Illustration of mouse PGC-1α (UniProt ID: O70343) with the CTD aligned against the human protein (UniProt ID: Q9UBK2). **b, c**, Heat maps with DEG (**b**) and predicted changes in TF activity (**c**) induced by overexpression of LacZ (control), PGC-1α FL or ΔCTD in C2C12 myotubes. **d, e**, Overlap (**d**) and gene ontology (GO) analysis (**e**) of core proteins regulated by PGC-1α FL or ΔCTD in C2C12 myotubes. Dashed line represents GO statistical cutoff (adjusted p-value < 0.05). **f**, Basal oxygen consumption rate (OCR) of C2C12 myotubes transduced with PGC-1α FL or ΔCTD at two different multiplicity of infections (MOI). Values are mean ± SD; **p < 0.01.

### PGC-1α function is impaired by deletion of the CTD

To investigate the function of the PGC-1α CTD, we overexpressed either full length (FL) PGC-1α or a truncated protein lacking the CTD (ΔCTD) in C2C12 myotubes. Both PGC-1α FL and ΔCTD exhibited comparable levels of overexpression at the RNA and protein level in a context of very low expression of endogenous PGC-1α (Supplementary Fig. 1a, b). Moreover, deletion of the CTD of PGC-1α did not affect protein half-life compared to FL protein (Supplementary Fig. 1c). In agreement with published data, transcriptome analysis revealed a large number of differentially expressed genes (DEG) upon overexpression of PGC-1α FL, whereas overexpression of ΔCTD induces a drastically different set of DEG (Fig. 1b, Supplementary Fig. 2a). Based on Gene Ontology (GO) analysis, the vast majority of DEG regulated by both PGC-1α FL and ΔCTD were related to metabolic pathways (Supplementary Fig. 2b), while the function of the ΔCTD-dependent DEG and those that are exclusively controlled by the CTD is less clearly defined (Supplementary Fig. 2b). We next leveraged our transcriptomic data to infer TFs regulated in a PGC-1α CTD-dependent manner by using Integrated System for Motif Activity Response Analysis (ISMARA)^10^. This analysis strongly suggests that the CTD of PGC-1α plays a central role in modulating the activity of a wide range of TFs (Fig. 1c). Interestingly, PGC-1α-induced activation of the nuclear receptor ERRα, known to bind at the N-terminal domain, was also predicted to be blunted in PGC-1α ΔCTD by ISMARA analysis, suggesting that CTD-specific features and mechanisms play a critical regulatory function.

The biological impact of the deletion of PGC-1α CTD was further investigated via whole proteome analysis of skeletal muscle cells. Proteome remodelling mediated by PGC-1α FL was reduced in PGC-1α ΔCTD, though the effect was milder compared to that observed at the transcriptome level (Supplementary Fig. 2c, d). Next, we defined a set of “core proteins” by integrating the transcriptomic and proteomic datasets, revealing genes transcriptionally up-or down-regulated by PGC-1α that exhibit a corresponding change at the protein level. We found that deletion of the CTD of PGC-1α drastically impaired the expression of such core proteins, which are primarily involved in transport processes and oxidative metabolism (Fig. 1d, e). Importantly, even though many core proteins in the overlap between PGC-1α FL and ΔCTD are also involved in oxidative phosphorylation and tricarboxylic acid cycle (Fig. 1e, Supplementary Fig. 2e, f), only overexpression of PGC-1α FL significantly increased basal oxygen consumption of skeletal muscle cells (Fig. 1f). Altogether, these data demonstrate that both the transcriptional and functional output of PGC-1α are highly modulated by the CTD.

### The PGC-1α CTD mediates RNA-dependent protein-protein interactions

Mechanistically, the CTD of PGC-1α represents a potential platform for protein-protein and protein-RNA interactions. By implementing an *in vitro* affinity purification followed by mass spectrometry (*in vitro* AP-MS) assay, we identified over 200 nuclear proteins interacting directly or indirectly with the CTD of PGC-1α (Fig. 2a). Most of PGC-1α CTD-interacting proteins were associated with the regulation of RNA processing and gene transcription (Fig. 2b). Consistently, analysis of our transcriptomic data demonstrated that PGC-1α FL regulates RNA processing events (Fig. 2c), corresponding to the alternative splicing of 203 transcripts (Fig. 2d). In contrast, over 60% of the alternative splicing capacity was lost when the CTD of PGC-1α was deleted (Fig. 2c, d). These data support and further expand previous findings proposing a role of PGC-1α in RNA processing via CTD-mediated protein-protein interactions^8,11^. It is however unknown to what extent direct RNA binding to the CTD of PGC-1α is involved in this process. We therefore carried out *in vitro* AP-MS in the absence and presence of RNase A, which revealed that 83% of PGC-1α CTD protein-protein interactions rely on RNA (Fig. 2e, Supplementary Fig. 3a, b). Many of the RNA-dependent interacting proteins are involved in skeletal muscle function, but a large number is involved in RNA processing, implying a direct link between the binding of RNAs and splicing factors to the CTD of PGC-1α (Fig. 2f, Supplementary Fig. 3c). Notably, RNA-dependent interactions also include several TFs (e.g. BTF3 and CEBPG) and components of transcriptional coregulator complexes (e.g. SGF29 and JMJD6) (Fig. 2f). These findings indicate that the CTD of PGC-1α regulates gene transcription and RNA processing in an RNA binding-dependent manner.

**Fig. 2:**
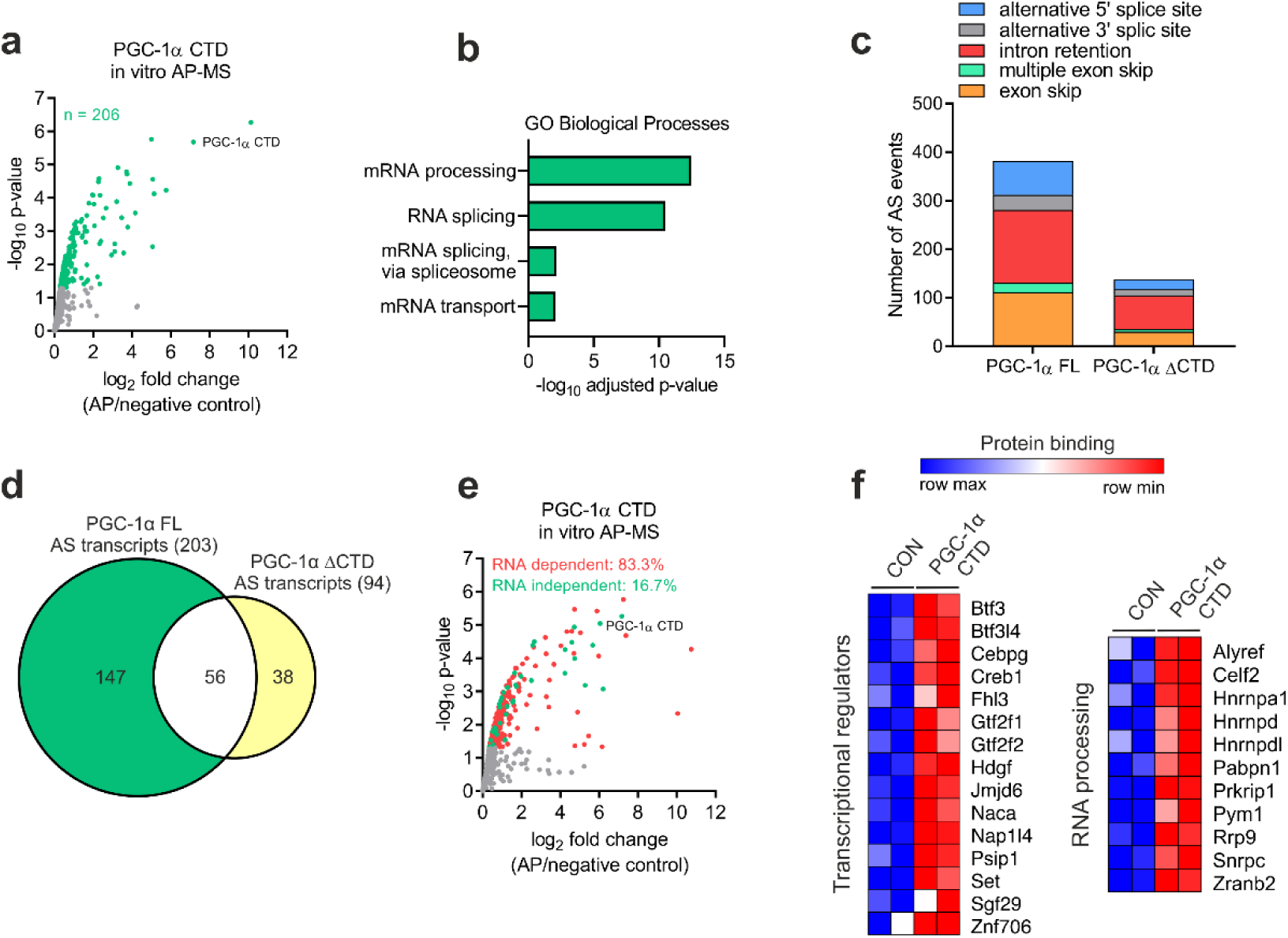
Most PGC-1α CTD protein interactions depend on RNA binding. **a**, Volcano plot with green dots representing proteins interacting with the CTD of PGC-1α. **b**, Gene ontology (GO) analysis of PGC-1α CTD interactome. **c, d**, Alternative splicing (AS) analysis of transcriptomic data showing the number of AS events (**c**) and transcripts (**d**) regulated by PGC-1α FL or ΔCTD in C2C12 myotubes. **e**, Volcano plot with red and green dots representing PGC-1α CTD RNA dependent and independent protein interactions, respectively. **f**, Heat maps showing PGC-1α CTD RNA dependent protein interactions with proteins regulating gene transcription and RNA processing.

### PGC-1α chromatin recruitment is regulated by RNA binding at the CTD

We next implemented *in vitro* cross-linking affinity purification followed by sequencing (*in vitro* uvAP-seq) to investigate the RNA-binding capacity of the PGC-1α CTD, which revealed direct interactions with a large number of RNAs (Fig. 3a). Interestingly, although PGC-1α can interact with protein-coding RNAs, the vast majority of bound RNAs were transcribed at regulatory elements such as promoters and distal intergenic regions (Fig. 3b, c), supporting and expanding previous reports of non-polyadenylated RNAs binding to PGC-1α^9^. Motif discovery analysis uncovered 11 motifs enriched among RNAs bound by the CTD (Fig. 3d). Such a diversity of RNA motifs has recently been found to be a common characteristic of a large number of RNA binding proteins (RBPs)^5,12^, including PGC-1α FL in primary hepatocytes^13^. Overall, our results now demonstrate that PGC-1α is a *bona fide* RBP, with a large promiscuity for binding several types of RNAs harbouring different recognition motifs.

**Fig. 3:**
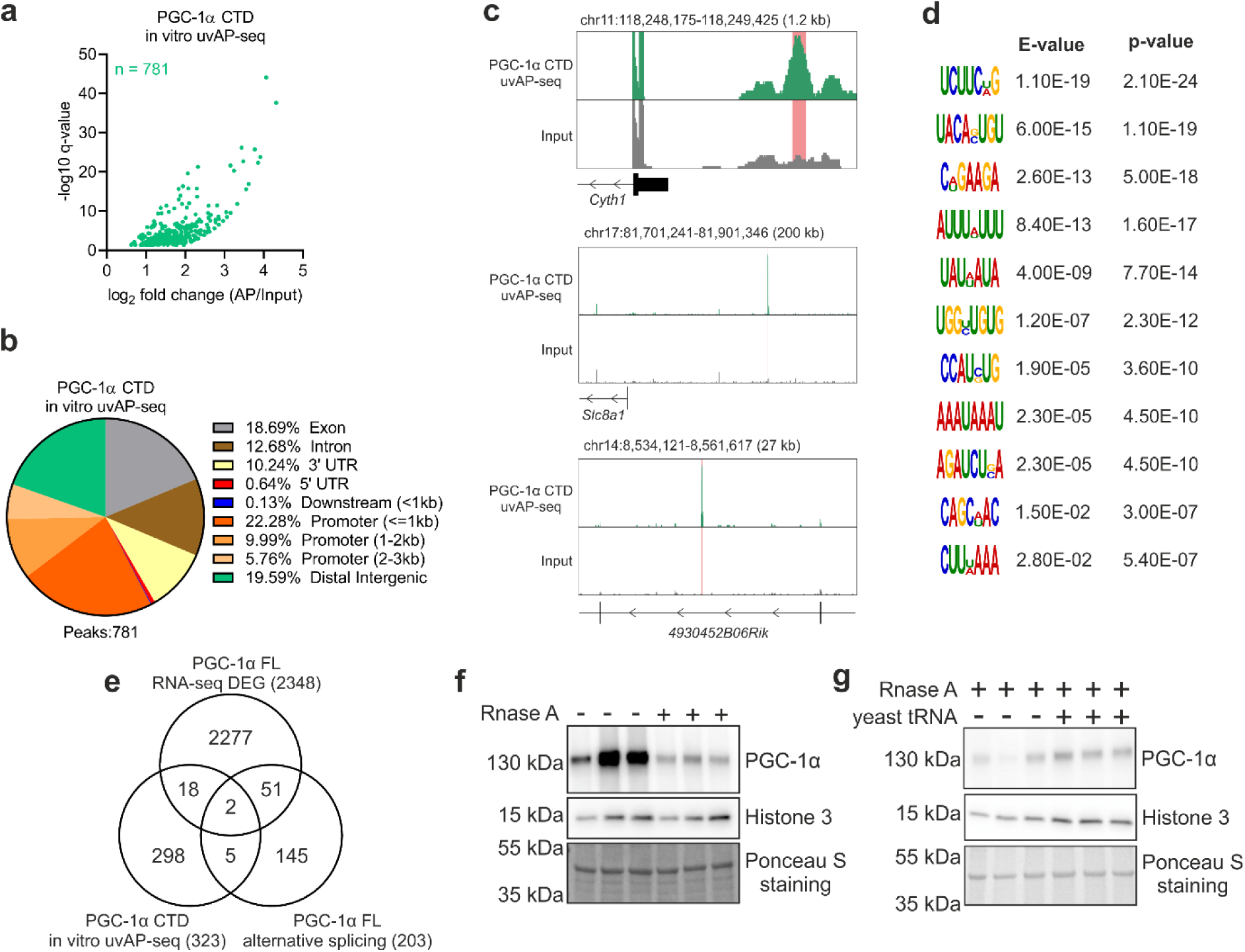
PGC-1α chromatin recruitment is regulated by RNA binding at the CTD. **a**, Volcano plot with green dots representing RNAs interacting with the CTD of PGC-1α. **b**, Annotation of *in vitro* AP-seq peaks representing PGC-1α CTD bound RNAs. **c**, Genome browser track view of representative peaks located at promoter (top panel), distal intergenic (middle panel) and intronic regions (bottom panel). **d**, Motif discovery analysis of PGC-1α CTD bound RNAs. **e**, Overlap of transcripts that are differentially expressed (DEG), alternative spliced and bound by PGC-1α. **f, g**, Protein content of PGC-1α FL bound to chromatin fractions in the absence and presence of 1 mg/ml of RNase A (**f**) or following the addition of 5 µg/µl of yeast tRNA (**g**). Immunoblots are representative of three independent experiments, each performed in triplicate. Ponceau S staining used as loading control in Fig. 3f was ran on a different gel.

Previous studies have linked PGC-1α CTD function to RNA processing^8^, e.g. for the stabilization of hepatic metabolic transcripts^13^. Integration of our RNA-seq and *in vitro* uvAP-seq datasets however revealed that over 90% of protein-coding RNAs bound by PGC-1α were neither DEG nor alternative spliced, while 97% of DEG were neither alternative spliced nor bound by PGC-1α (Fig. 3e). While the small overlap between RNA binding and differential gene expression was also observed in the liver^13^, our results indicate that RNA processing is not the main function of RNA binding to the CTD. Chromatin-bound RBPs have recently emerged as key regulators of gene transcription at active promoters and enhancers^12,14^. Moreover, PGC-1α activity can be modulated by binding of enhancer and other long noncoding RNAs at DNA regulatory elements^9,15^. A comparison of our RNA binding and PGC-1α FL ChIP-seq data^16^ demonstrates a low correlation between RNA-and chromatin-binding profiles (Supplementary Fig. 4a, b), similar to what has been reported for many other RBPs^12,14^. Thus, RNA interactions with the CTD of PGC-1α at the site of genomic recruitment, e.g the reported interaction with enhancer RNAs^9^, seems to be a minor event. Therefore, to define how RNA binding modulates PGC-1α function, we measured its recruitment to chromatin in the absence and presence of RNase A via subcellular fractionation. This experiment revealed that the absence of nuclear RNAs strongly decreases the amount of chromatin-bound PGC-1α FL (Fig. 3f). Importantly, subsequent reloading of nuclei with yeast tRNA was sufficient to restore chromatin-bound PGC-1α FL levels (Fig. 3g). Our data therefore demonstrate that, besides regulating protein-protein interactions, RNA binding at the CTD of PGC-1α is a crucial mechanism controlling its dynamic recruitment to chromatin.

### The CTD of PGC-1α regulates sub-nuclear localization at transcriptional condensates

While residual levels of PGC-1α FL in cytoplasm and nucleoplasm of C2C12 myotubes were observed, we found that most of the protein was strongly bound to chromatin (Fig. 4a). In stark contrast, chromatin binding of PGC-1α ΔCTD was markedly reduced, associated with a shift towards higher relative levels in the nucleoplasm and cytoplasm compared to the FL protein (Fig. 4a). Intriguingly, both endogenous and GFP-fused PGC-1α FL proteins form characteristic nuclear foci, originally defined as nuclear speckles^8,15^. We now observed that deletion of the CTD of GFP-fused PGC-1α completely abolished its localization in nuclear foci in C2C12 myoblasts (Fig. 4b). Together, our data indicate that a large fraction of PGC-1α foci represent a chromatin-rich sub-compartment such as chromatin condensates, rather than nuclear speckles that are thought to reside in the nucleoplasm^17^.

**Fig. 4:**
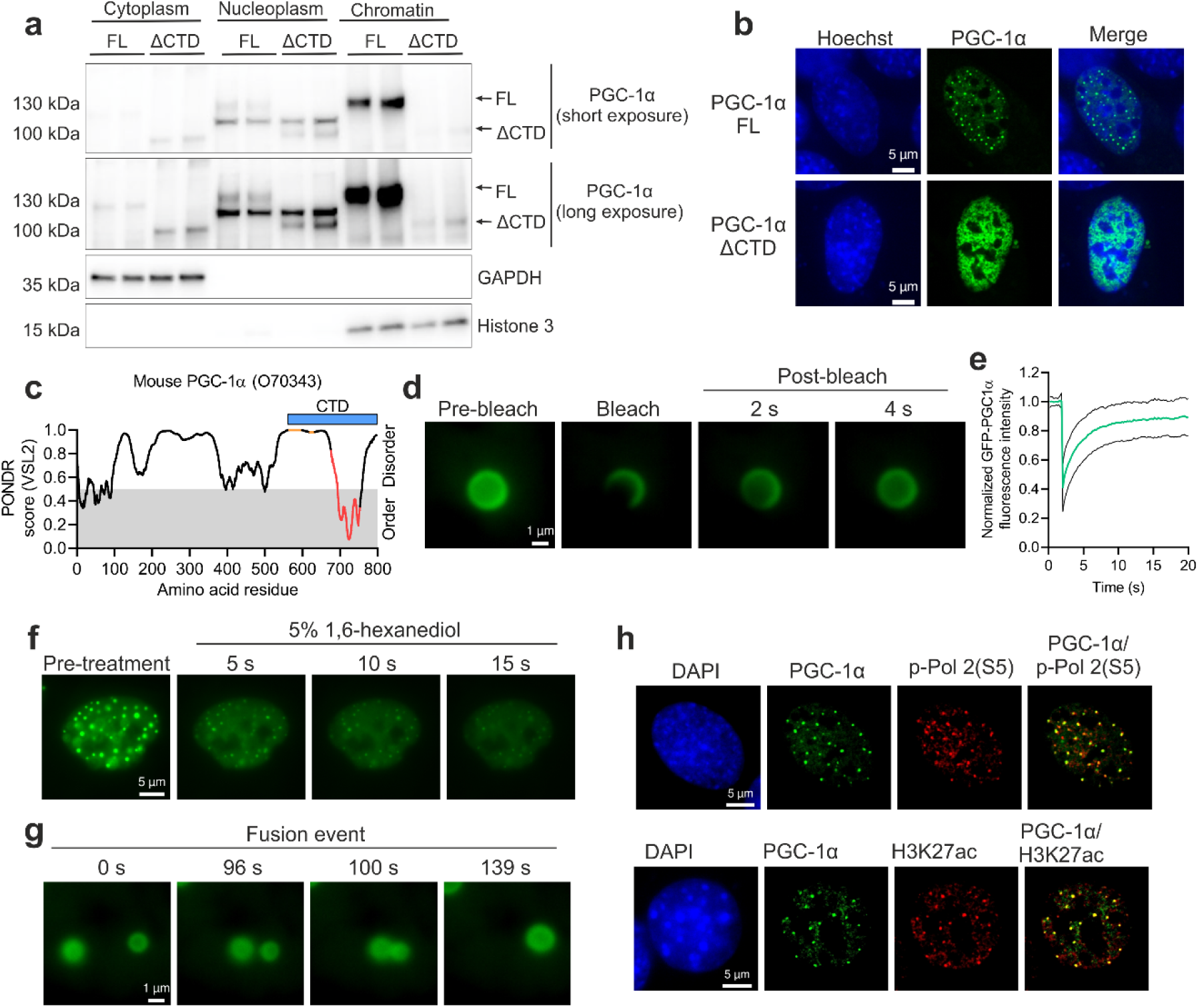
PGC-1α activates gene transcription within liquid-like condensates. **a**, Subcellular fractionation of C2C12 myotubes expressing PGC-1α FL or ΔCTD. **b**, Nuclear localization of GFP-PGC-1α FL or ΔCTD fusion proteins transfected in C2C12 myoblasts. **c**, Predictor of Natural Disordered Regions (PONDR) analysis of mouse PGC-1α protein, with orange dots, red dots and blue bar representing RS domains, RRM and the CTD region under investigation, respectively. **d, e**, Live-cell imaging (**d**) and quantification (**e**) of GFP-PGC-1α FL FRAP experiments in C2C12 myoblasts. Green and black lines denote mean and SD of FRAP quantification, respectively. **f**, Live-cell imaging of 5% 1,6-hexanediol treatment of C2C12 myoblasts transfected with GFP-PGC-1α FL. **g**, Live-cell imaging of a GFP-PGC-1α FL droplet fusion event in C2C12 myoblasts. **h**, Images of GFP-PGC-1α FL co-stained with DAPI (DNA), p-Pol 2(S5) and H3K27ac in C2C12 myoblasts. Immunoblots and microscopy images are representative of at least three independent experiments each in triplicate.

Chromatin condensates are membrane-less organelles formed via liquid-liquid phase separation (LLPS) that play a critical regulatory role in gene expression^18,19^. Interestingly, the nuclear foci containing PGC-1α resemble nuclear condensates formed by TFs and transcriptional coregulators^20,21^. The formation of liquid-like droplets is often driven by the presence of intrinsically disordered regions (IDR) on proteins and by multivalent protein and RNA interactions of RBPs^22^. PGC-1α is largely disordered (Fig. 4c, Supplementary Fig. 5a), yet our findings demonstrate that N-terminal IDRs alone are insufficient to mediate the formation of nuclear foci (Fig. 4b). Conversely, in presence of the RRM, PGC-1α is found in foci (Fig. 4b), implying that RNA binding plays an important role in localising PGC-1α within nuclear foci. To assess potential liquid-like properties of PGC-1α foci, we carried out fluorescence recovery after photobleaching (FRAP) analysis that revealed a rapid fluorescence recovery characteristic of liquid-like condensates (Fig. 4d, e). We additionally studied the physical properties of PGC-1α foci by using the aliphatic alcohol 1,6-hexanediol, which resulted in a dissolving of foci within seconds (Fig. 4f, Supplementary Fig. 5b). Finally, the liquid-like properties of PGC-1α-containing condensates was supported by live cell imaging-based detection of fusion events between different foci (Fig. 4g, Supplementary Fig. 5c). While the precise molecular composition of these condensates awaits clarification, it is tempting to speculate that as a strong transcriptional coactivator, PGC-1α might represent a sub-compartment favouring active gene transcription by assembling transcriptional multi-protein complexes. Consistent with this hypothesis, we found that PGC-1α condensates exhibit an enrichment of Pol 2 phosphorylated at S5 of its CTD (p-Pol 2(S5)) and histone 3 acetylated at K27 (H3K27ac), thus reflecting active transcription at enhancers and promoter (Fig. 4h). Therefore, our findings demonstrate that PGC-1α condensates compartmentalize the transcriptional machinery to concentrate and stimulate gene transcription at DNA regulatory elements.

Collectively, our results reveal key novel insights into the regulation of complex transcriptional networks, where coregulators have been heralded as master controllers using so far elusive mechanisms^23,24^. By studying PGC-1α as a prototypical member of the coactivator protein superfamily, we now demonstrate that its highly conserved CTD fulfils a crucial role for controlling gene expression by compartmentalizing multi-protein complexes of transcriptional regulators within nuclear condensates. Remarkably, the assembly of such multi-protein complexes on the chromatin is dynamically regulated by PGC-1α-RNA binding, which might facilitate the spatiotemporal adaptive response to stimuli such as exercise and diet^25^. These unprecedented mechanistic insights now outline how transcriptional coregulators could interpret environmental cues to control transcriptional networks in a context-specific manner via assembly of nuclear condensates containing multi-protein transcriptional complexes and specific DNA regulatory elements.

Physical inactivity, which impairs skeletal muscle metabolism and function, can accelerate the development of a wide spectrum of diseases, including type 2 diabetes, cancer and non-alcoholic fatty liver disease^26^. Modulation of PGC-1α nuclear condensate formation thus represents an attractive strategy to regulate the function of this transcriptional coactivator. Notably, the formation of abnormal liquid-like condensates has recently been found to be associated with the development of pathologies such as cancer and neurodegenerative diseases^27^. Hence, the fundamental mechanism controlling PGC-1α function discovered in this work might have important implication in the development of strategies to improve energy metabolism under pathological conditions, e.g. by leveraging pharmacological approaches aimed at the selective modulation of transcriptional condensates^28^.

## Supplementary figures

**Supplementary Fig. 1:**
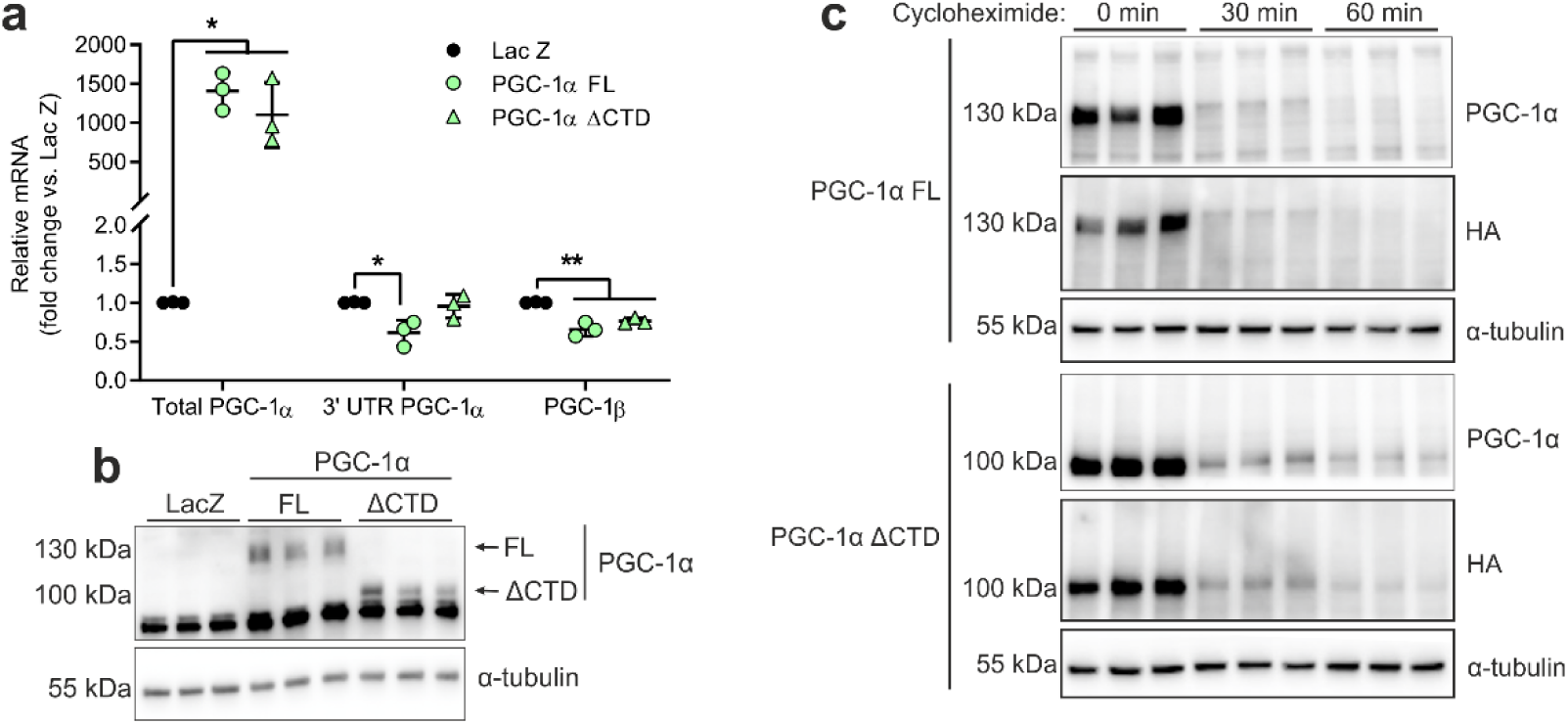
Overexpression of PGC-1α FL and ΔCTD in C2C12 myotubes. **a, b**, PGC-1α mRNA (**a**) and protein (**b**) levels in C2C12 myotubes transduced with LacZ (control), PGC-1α FL or ΔCTD (n = 3 independent experiments each in triplicate). **c**, PGC-1α protein half-life analysis in C2C12 myotubes transduced with PGC-1α FL or ΔCTD and treated with DMSO as control or 100 μg/ml cycloheximide. Values are mean ± SD; *p < 0.05 and **p < 0.01. Immunoblots are representative of three independent experiments, each performed in triplicate.

**Supplementary Fig. 2:**
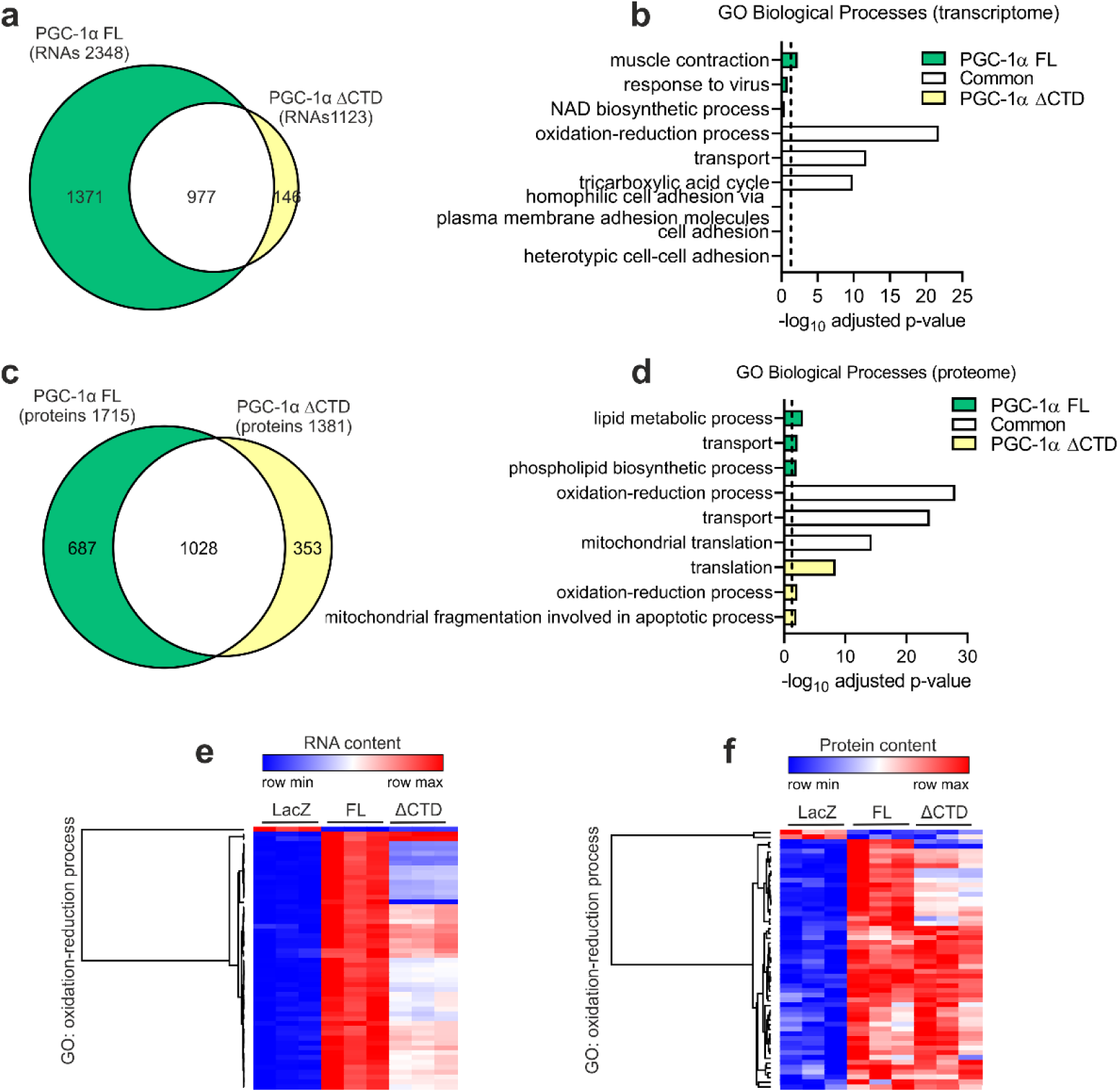
Transcriptome and proteome analysis. **a, b**, Overlap (**a**) and gene ontology (GO) analysis (**b**) of DEG induced by PGC-1α FL or ΔCTD in C2C12 myotubes. **c, d**, Overlap (**c**) and GO analysis (**d**) of proteins regulated by PGC-1α FL or ΔCTD in C2C12 myotubes. **e, f**, Heat maps showing the content of RNAs (**e**) and proteins (**f**) contained in the GO term oxidation-reduction process in C2C12 myotubes overexpressing LacZ (control), PGC-1α FL or ΔCTD. Dashed line represents GO statistical cutoff (adjusted p-value < 0.05).

**Supplementary Fig. 3:**
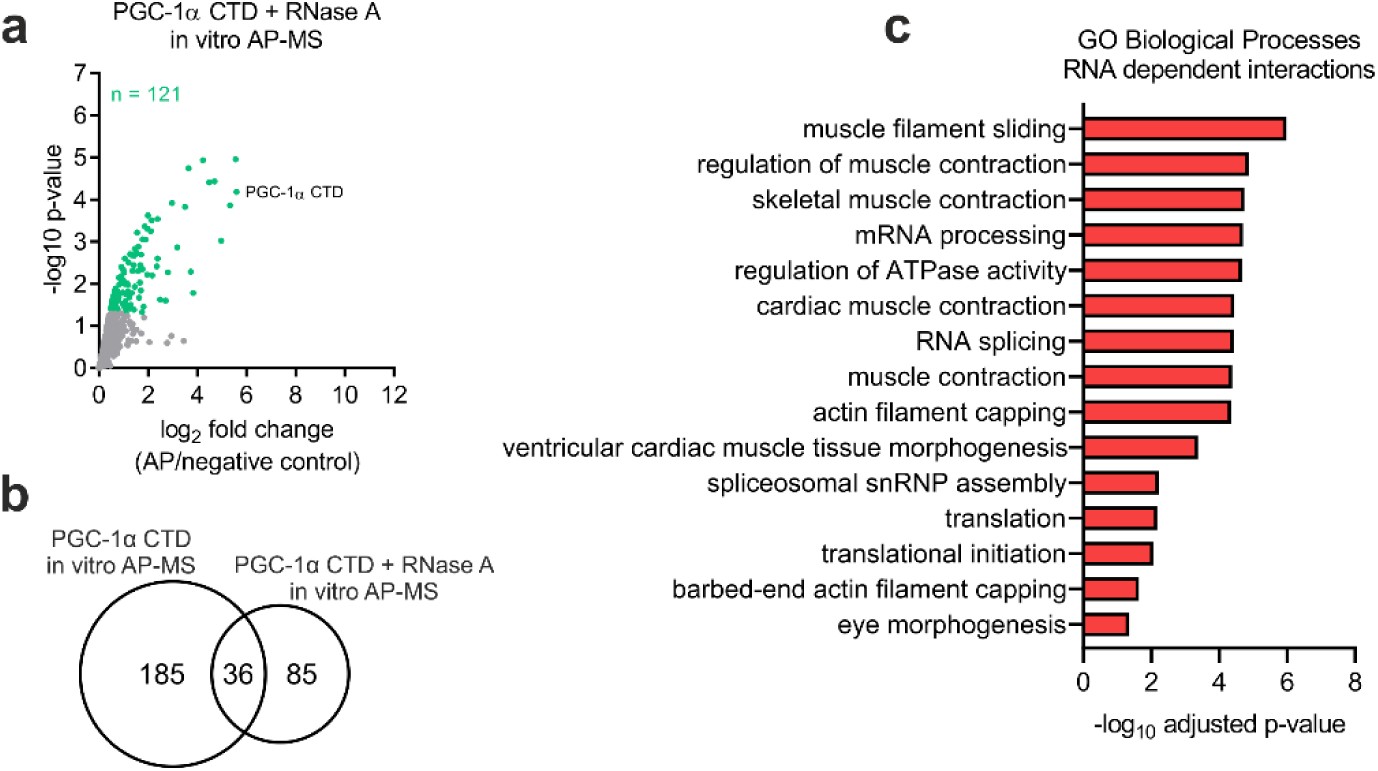
RNA dependent protein-protein interaction analysis. **a**, Volcano plot with green dots representing proteins interacting with the CTD of PGC-1α in the presence of 1 mg/ml of RNase A. **b**, Overlap of PGC-1α CTD interacting proteins in the absence and presence of 1 mg/ml of RNase A. **c**, Gene ontology (GO) analysis of RNA dependent PGC-1α CTD interacting proteins.

**Supplementary Fig. 4:**
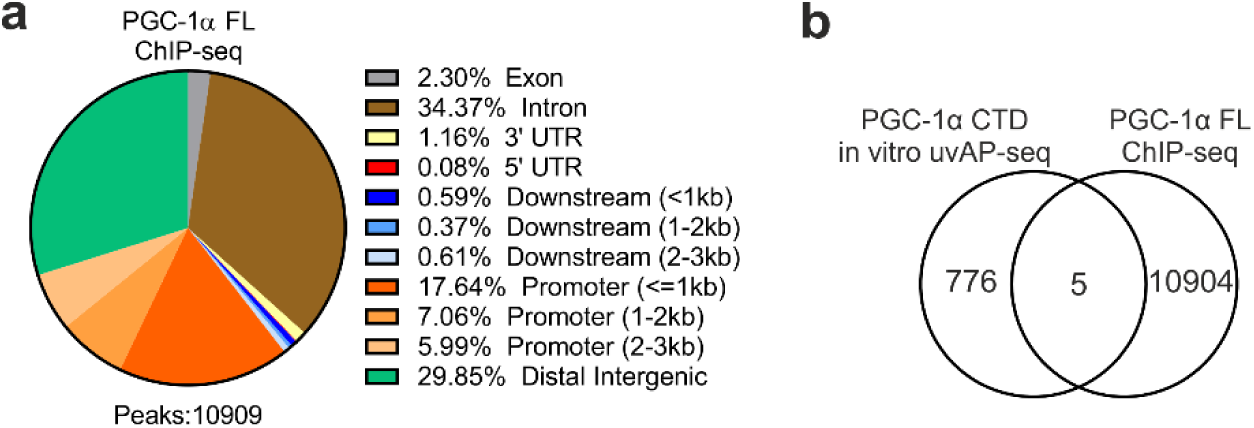
ChIP-seq analysis of PGC-1α FL. **a**, Annotation of PGC-1α FL ChIP-seq peaks. **b**, Overlap of PGC-1α CTD and FL in vitro AP-seq and ChIP-seq peaks, respectively.

**Supplementary Fig. 5:**
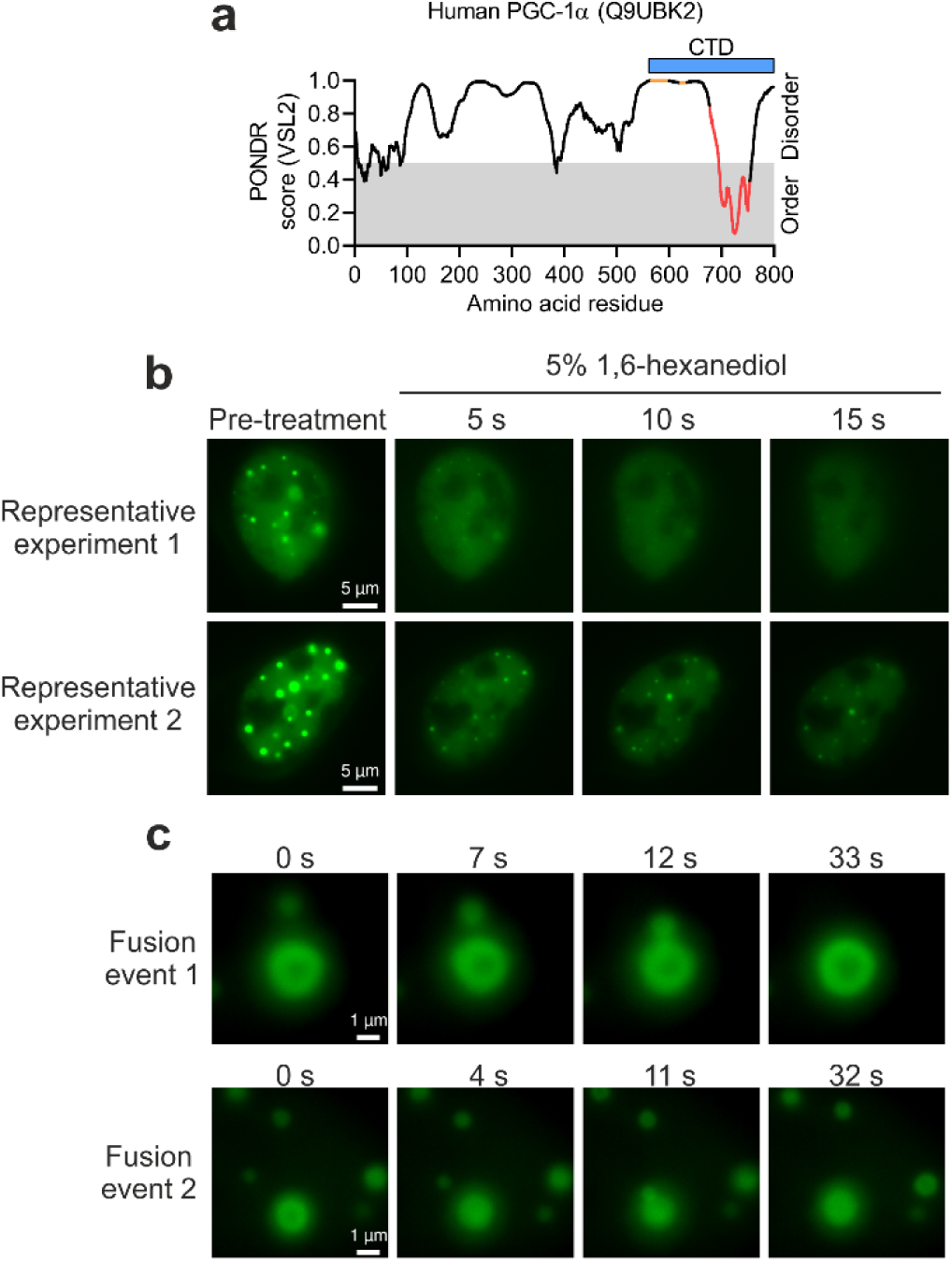
Analysis of liquid-like properties of PGC-1α nuclear foci. **a**, Predictor of Natural Disordered Regions (PONDR) analysis of human PGC-1α protein, with orange and red dots representing RS domains and RRM, respectively. **b**, Additional live imaging of 5% 1,6-hexanediol treatment of C2C12 myoblasts transfected with GFP-PGC-1α FL. **c**, Additional live imaging of GFP-PGC-1α FL droplet fusion events in C2C12 myoblasts. Microscopy images are representative of at least three independent experiments, each performed in triplicate.

## Methods

### Cell culture

C2C12 myoblasts were grown in Dulbecco’s modified Eagle’s medium (DMEM) supplemented with 10% fetal bovine serum (growth medium). To induce differentiation, growth medium of about 90% confluent myoblasts was changed to DMEM supplemented with 2% horse serum (differentiation medium). Experiments using C2C12 myotubes were performed after 4 days of differentiation. Cells were maintained at 37°C, 95% O_2_ and 5% CO^2^.

### Generation of DNA plasmids and transfections

Mouse GFP-PGC-1α full length (FL) plasmid (Addgene, #4) was used as template for PCR amplification of a delta C-terminal domain (ΔCTD) fragment containing amino acid 1-564 of PGC-1α. SalI and BamHI restriction sites were added to the 5’ and 3’ end of the insert, respectively. The destination plasmid GFP-PGC-1α FL was digested with SalI and BamHI, following which the PGC-1α ΔCTD insert was ligated. Plasmid was corroborated via Sanger sequencing.

Transfection of C2C12 myoblasts was performed using Opti-MEM(tm) (Thermo Fisher Scientific, #31985070) and Lipofectamine(tm) 2000 (Thermo Fisher Scientific, #11668019) following manufacturer’s instructions. Twenty four hours after seeding, cells were transfected with 0.5 μg of EGFP-PGC-1α FL (Addgene, #4) or ΔCTD for 24 h.

### Generation of adenoviral vectors

Adenovirus vectors were generated with the Adeno-X Adenoviral System 3 following manufacturer’s instructions (Takara, #632267). Briefly, mouse PGC-1α FL and ΔCTD were PCR-amplified from the pcDNA-f:PGC1 (Addgene, # 1026) and pcDNA-f:PGC1(delta CTD) (Addgene, #1030) plasmid vectors, respectively. N-terminal HA and FLAG tags were introduced during PCR amplification, with the amplicon subcloned into the pAdenoX-ZsGreen1 vector to generate HA-Flag-PGC-1α FL and ΔCTD adenoviruses. The *LacZ* gene was also subcloned into the pAdenoX-ZsGreen1 vector, which was used to generate a LacZ control adenovirus. All plasmids were corroborated via Sanger sequencing. Adenoviruses were produced and amplified in Adeno-X(tm) 293 cells (Takara, # 632271), while titter was determined by fluorescence-activated cell sorting.

### Adenovirus transduction

Cells were transduced with HA-Flag-PGC-1α FL, ΔCTD or LacZ adenovirus at multiplicity of infection (MOI) 2-6. Adenoviruses were prepared in the corresponding medium and cells were transduced for 4 h. Next, cells were washed once with phosphate buffered saline (PBS) and, then, incubated in adenovirus-free medium for a total of 48 h.

### RNA purification and quantitative PCR (qPCR)

Cells were collected in TRI Reagent (Sigma #T9424), following which RNA was purified and reverse transcribed using Direct-zol(tm) RNA MiniPrep (Zymo Research, #R2050) and iScript(tm) cDNA Synthesis Kit (Bio-Rad, #1708891), respectively. Relative changes in mRNA content was quantified by qPCR on a StepOnePlus system (Applied Biosystems) using Fast SYBR(tm) Green Master Mix (Thermo Fisher Scientific, #4385612). The ΔΔCT method was used for analysis, with TATA binding protein (*Tbp*) as endogenous control.

### RNA sequencing (RNA-seq) and alterative splicing analysis

Libraries were prepared with TruSeq Stranded mRNA Library Kit (Illumina, #20020595), pair-end sequencing was performed using the HiSeq 2500 (Illumina) and data was analysed on the Galaxy platform (https://usegalaxy.eu/). Reads were trimmed with Trim Galore! (Galaxy version 0.4.3.1) and quality was assessed using FastQC (Galaxy version 0.72+galaxy1). Reads were aligned to the mm10 version of the mouse genome using STAR (Galaxy version 2.7.2b), while strand specificity and read counting was performed with Infer Experiment (Galaxy version 2.6.4.1) and featureCounts (Galaxy version 1.6.4+galaxy1), respectively. Next, we used DESeq2 (Galaxy version 2.11.40.6) for differential expression analysis (q-value < 0.01 and fold change ≥ 2) and the resulting data was annotated with Annotate DESeq2/DEXSeq output tables (Galaxy version 1.1.0). Overlap between different datasets was determined with Venny (version 2.1, https://bioinfogp.cnb.csic.es/tools/venny/). Gene Ontology (GO) analysis was performed with DAVID 6.8 (https://david.ncifcrf.gov/), with significance defined as adjusted (Benjamini) p-value < 0.05. Transcription factors activity analysis was achieved with ISMARA (https://ismara.unibas.ch/mara/), where z-value > 2 was consider as significant. We used DESeq2 normalized counts to generate heat maps and for hierarchical clustering using Morpheus (https://clue.io/morpheus). Finally, in order to identify alternative splicing events the SplAdder workflow was employed^29^. We produced splicing graphs for all samples belonging to a comparison we were interested in. Afterwards we called splicing events (exon_skip, intron_retention, alt_3prime, alt_5prime, mult_exon_skip) with the SplAdder test procedure using default parameters.

### Protein extraction and immunoblotting

Protein was extracted with protein lysis buffer (50 mM Tris-HCl (pH 7.5), 150 mM NaCl, 1 mM EDTA (pH 8.0), 5 % Glycerol, 1 % NP40, 0.1 % SDS, 1 mM nicotinamide, 1X Halt(tm) Protease Inhibitor Cocktail (Thermo Fisher Scientific, #87786)) as previously described^30^.

Immunoblotting was performed with precast (Bio-Rad, #4561096) or home-made SDS-polyacrylamide gels as previously described^30^, and proteins were detected with a primary antibody to PGC-1α (Santa Cruz Biotechnology, #sc-518025), α-tubulin (Cell Signaling Technology, #2144S), histone 3 (Abcam, #ab1791), GAPDH (Cell Signaling Technology, #2118S) and HA (Sigma, #11867423001). If required, Ponceau S (Sigma, #P7170-1L) staining was used as loading control. Secondary antibodies for mouse (Agilent, #P0260), rabbit (Agilent, #P0399) and rat (Jackson ImmunoResearch, #112-035-003) were used. Antibody binding was detected using enhanced chemiluminescence HRP substrate detection kit for standard (Thermo Fisher Scientific, #32106), medium (Thermo Fisher Scientific, #34076) or high (Thermo Fisher Scientific, #34095) sensitivity.

### Protein half-life

Protein half-life was measured by treating cells with DMSO as control or 100 μg/ml of cycloheximide (Sigma, #C4859) for 30 or 60 min. Next, cells were collected for protein extraction and immunoblotting.

### Mass spectrometry analysis of whole cell proteome

Cells were lysed in 80 μl of lysis buffer (1% sodium deoxycholate (SDC), 0.1 M TRIS, 10 mM TCEP, pH = 8.5) using 10 cycles of sonication (Bioruptor, Diagnode). Samples were reduced for 10 min at 95°C and alkylated at 15 mM chloroacetamide for 30 min at 37°C. Proteins were digested by incubation with sequencing-grade modified trypsin (1/50 w/w; Promega,V5113) for 12 h at 37°C. Tryptic digests were acidified (pH<3) using TFA and cleaned up using iST cartridges (PreOmics, P.O.00027) according to the manufacturer’s instructions. Samples were dried under vacuum and stored at −20 °C.

Sample aliquots comprising 25 μg of peptides were labelled with isobaric tandem mass tags (TMT 10-plex, Thermo Fisher Scientific, 90110) as described previously^31^. Shortly, peptides were re-suspended in 20 μl labelling buffer (2 M urea, 0.2 M HEPES, pH 8.3) and 5 μL of each TMT reagent were added to the individual peptide samples followed by a 1 h incubation at 25°C, shaking at 500 rpm. To quench the labelling reaction, 1.5 μL aqueous 1.5 M hydroxylamine solution was added and samples were incubated for another 10 min at 25°C shaking at 500 rpm followed by pooling of all samples. The pH of the sample pool was increased to 11.9 by adding 1 M phosphate buffer (pH 12) and incubated for 20 min at 25°C shaking at 500 rpm to remove TMT labels linked to peptide hydroxyl groups. Subsequently, the reaction was stopped by adding 2 M hydrochloric acid until a pH < 2 was reached. Finally, peptide samples were further acidified using 5 % TFA, desalted using Sep-Pak Vac 1cc (50 mg) C18 cartridges (Waters, WAT054960) according to the manufacturer’s instructions and dried under vacuum.

TMT-labelled peptides were fractionated by high-pH reversed phase separation using a XBridge Peptide BEH C18 column (3,5 µm, 130 Å, 1 mm x 150 mm; Waters, 186003562) on an Agilent 1260 Infinity HPLC system. Peptides were loaded on column in buffer A (20 mM ammonium formate in water, pH 10) and eluted using a two-step linear gradient from 2% to 10% in 5 minutes and then to 50% buffer B (20 mM ammonium formate in 90% acetonitrile, pH 10) over 55 minutes at a flow rate of 42 µl/min. Elution of peptides was monitored with a UV detector (215 nm, 254 nm) and a total of 36 fractions were collected, pooled into 12 fractions using a post-concatenation strategy as previously described^32^ and dried under vacuum.

Dried peptides were re-suspended in 0.1% aqueous formic acid and subjected to LC–MS/MS analysis using a Q Exactive HF Mass Spectrometer fitted with an EASY-nLC 1000 (Thermo Fisher Scientific) and a custom-made column heater set to 60°C. Peptides were resolved using a RP-HPLC column (75μm × 30cm) packed in-house with C18 resin (ReproSil-Pur C18–AQ, 1.9 μm resin; Dr. Maisch, r119.aq.) at a flow rate of 0.2 μLmin-1. The following gradient was used for peptide separation: from 5% B to 15% B over 10 min to 30% B over 60 min to 45 % B over 20 min to 95% B over 2 min followed by 18 min at 95% B. Buffer A was 0.1% formic acid in water and buffer B was 80% acetonitrile, 0.1% formic acid in water.

The mass spectrometer was operated in DDA mode with a total cycle time of approximately 1 s. Each MS1 scan was followed by high-collision-dissociation (HCD) of the 10 most abundant precursor ions with dynamic exclusion set to 30 s. For MS1, 3e6 ions were accumulated in the Orbitrap over a maximum time of 100 ms and scanned at a resolution of 120,000 FWHM (at 200 m/z). MS2 scans were acquired at a target setting of 1e5 ions, maximum accumulation time of 100 ms and a resolution of 30,000 FWHM (at 200 m/z). Singly charged ions and ions with unassigned charge state were excluded from triggering MS2 events. The normalized collision energy was set to 35%, the mass isolation window was set to 1.1 m/z and one microscan was acquired for each spectrum.

The acquired raw-files were converted to the mascot generic file (mgf) format using the msconvert tool (part of ProteoWizard, version 3.0.4624 (2013-6-3)) and searched using MASCOT against a murine database (consisting of 49434 forward and reverse protein sequences downloaded from Uniprot on 20141124) and 390 commonly observed contaminants. The precursor ion tolerance was set to 10 ppm and fragment ion tolerance was set to 0.02 Da. The search criteria were set as follows: full tryptic specificity was required (cleavage after lysine or arginine residues unless followed by proline), 3 missed cleavages were allowed, carbamidomethylation (C) and TMT6plex (K and peptide N-terminus) were set as fixed modification and oxidation (M) as a variable modification. Next, the database search results were imported into the Scaffold Q+ software (version 4.3.2, Proteome Software Inc.) and the protein false discovery rate was set to 1% based on the number of decoy hits. Proteins that contained similar peptides and could not be differentiated based on MS/MS analysis alone were grouped to satisfy the principles of parsimony. Proteins sharing significant peptide evidence were grouped into clusters. Acquired reporter ion intensities in the experiments were employed for automated quantification and statistical analysis using a modified version of our in-house developed SafeQuant R script v2.3^31^. This analysis included adjustment of reporter ion intensities, global data normalization by equalizing the total reporter ion intensity across all channels, summation of reporter ion intensities per protein and channel, calculation of protein abundance ratios and testing for differential abundance using empirical Bayes moderated t-statistics. The calculated p-values were corrected for multiple testing using the Benjamini–Hochberg method, with significance defined as q-value < 0.05 and fold change ≥ 1.2. Overlap between different datasets was determined with Venny (version 2.1, https://bioinfogp.cnb.csic.es/tools/venny/). GO analysis was performed with DAVID 6.8 (https://david.ncifcrf.gov/), with significance defined as adjusted (Benjamini) p-value < 0.05.

### Assessment of oxygen consumption

Basal oxygen consumption was measured using the Seahorse XF Cell Mito Stress Test Kit (Agilent, #103015-100) on a Seahorse XF96 Analyzer (Agilent) according to the manufacturer’s instructions.

### Subcellular fractionation

Subcellular fractionation was performed as previously described^33,34^, with the following modifications. Cells, on 10 cm plates, were washed twice with ice-cold PBS and collected into 5 ml of ice-cold PBS. Samples were centrifuged for 2 min at 500 g at 4°C and the cell pellet was re-suspended in 500 μl of ice-cold cytoplasmic lysis buffer (0.15% NP-40, 10 mM Tris-HCl (pH 7.0), 150 mM NaCl and 1X Halt(tm) Protease Inhibitor Cocktail (Thermo Fisher Scientific, #87786)). Following 5 min incubation on ice, samples were homogenized by using a glass Donce homogenizer with 20 strokes with a tight pestle on ice. The resulting cell lysate was layered onto 1250 μl of ice-cold sucrose buffer (10 mM Tris-HCl (pH 7.0), 150 mM NaCl, 25% sucrose and 1X Halt(tm) Protease Inhibitor Cocktail) and centrifuged for 10 min at 16000 g at 4°C. The supernatant containing the cytoplasmic fraction was transferred to a pre-chilled tube and snap frozen in liquid nitrogen. Nuclei pellet was wash once with 1 ml of nuclei wash buffer (0.1% Triton X-100, 1mM EDTA and 1X Halt(tm) Protease Inhibitor Cocktail in PBS) and centrifuged for 1 min at 1150 g at 4°C. Nuclei pellet was re-suspended in 200 μl of glycerol buffer (20 mM Tris-HCl (pH 8.0), 75 mM NaCl, 0.5 mM EDTA, 50% glycerol, 0.85 mM DTT and 1X Halt(tm) Protease Inhibitor Cocktail), after which 200 μl of nuclei lysis buffer (1% NP-40, 20 mM HEPES (pH 7.5), 300 mM NaCl, 1M urea, 0.2 mM EDTA, 1 mM DTT and 1X Halt(tm) Protease Inhibitor Cocktail) was added. Samples were mixed by pulsed vortex and incubate on ice for 2 min. Following centrifugation for 2 min at 18500 g at 4°C, the supernatant containing the nucleoplasm fraction was transferred to a pre-chilled tube and snap frozen in liquid nitrogen. The chromatin pellet was incubated with 50 μl of 1X RQ1 DNase Reaction Buffer with 30U of RQ1 RNase-Free DNase (Promega, # M6101) for 10 min at 37°C. Next, samples were placed on ice and 50 μl of storage buffer (10 mM Tris (pH 7.4), 1 mM EDTA, 25 mM NaCl, 10% glycerol and 1X Halt(tm) Protease Inhibitor Cocktail) were added, following which sonication was performed using Bioruptor® Plus (Diagenode) at 4°C for 15 min with 30 sec on and 30 sec off. Finally, samples were centrifuged at 16100 g for 10 min at 4 °C and the supernatant containing the chromatin was transferred to a pre-chilled tube and snap frozen in liquid nitrogen. Samples were prepared for immunoblotting as described above.

### RNase A and yeast tRNA treatment of nuclei

Nuclei were isolated as described above, washed once with 1 ml of ice-cold PBS and centrifuged for 1 min at 1150 g at 4°C. The pellet was re-suspended in 500 μl of 0.05% Tween-20 in PBS and incubated on ice for 10 min to permeabilize the nuclei. Next, samples were centrifuged for 1 min at 1150 g at 4°C and washed once with 1 ml of ice-cold PBS. Samples were re-suspended in either 100 μl of PBS with or without 1 mg/ml of RNase A (Sigma, #R4642) and were incubated for 15 min at 37°C. Next, samples were centrifuged at 2300 g for 10 min at 4°C and chromatin fraction was extracted as described above.

For tRNA treatments all samples were digested with RNase A, following which samples were washed twice with 1 ml of ice-cold PBS. Nuclei pellets were re-suspended in 100 µl of PBS containing 1 U/μl RNasin® Ribonuclease Inhibitors (Promega, # N2615) with or without 5 µg/µl of yeast tRNA (Thermo, #AM7119) and were then incubated for 15 min at 37°C. Subsequently, 1 ml of nuclei wash buffer (0.1% Triton X-100, 1mM EDTA and 1X Halt(tm) Protease Inhibitor Cocktail in PBS) was added and samples were centrifuged for 1 min at 1150 g at 4°C. Chromatin fraction was extracted as described above.

### In vitro affinity purification mass spectrometry (in vitro AP-MS)

C2C12 myotubes were transduced with PGC-1α FL at MOI 3 and nuclear extract was prepared with NE-PER(tm) Nuclear and Cytoplasmic Extraction Reagents (Thermo Fisher Scientific, #78835). Pull-down and negative control comprised 350 μg of protein from nuclear extract, while 12 μg of recombinant N-terminal His-Tag CTD of mouse PGC-1α (US Biological Life Sciences, #156296) was added only to pull-down samples. Samples were incubated overnight at 4°C with rotation, following which protein-protein complexes were purified with Pierce(tm) His Protein Interaction Pull-Down Kit (Thermo Fisher Scientific, #21277) following manufacturer’s instructions. RNA-dependent interactions were assessed by treating nuclear extracts with 1 U/μl RNasin® Ribonuclease Inhibitors (Promega, # N2615) as control or 1 mg/ml of RNase A (Sigma, #R4642) for 15 min at 37°C before adding the recombinant protein for overnight incubation.

Eluted proteins were incubated with four volumes of 100% trichloroacetic acid on ice for 10 min. Samples were then centrifuged at 18500 g for 5 min and the protein pellet was washed twice with 200 μl of cold acetone. The final protein pellet was re-suspended in 40 ul Gua buffer (2 M Guanidinium-HCl, 0.1 M Ammonium bicarbonate and 5 mM TCEP), sonicated ten times with Vial Tweeter ultrasonicator (Hielscher) and incubated for 10 min at 95°C, followed by alkylation of proteins with 15 mM chloroacetamide for 30 min at 37°C. Next, guanidium-HCl was diluted below 0.4 M with 0.1 M ammonium bicarbonate prior adding 0.5 ug trypsin and incubated for 12 h at 37°C shaking at 300 rpm. Tryptic digest was acidified (pH<3) using TFA and desalted using C18 reverse phase spin columns (Microspin, The Nest Group, Inc., #SEM SS18V) according to the manufacturer’s instructions. Peptides were dried under vacuum and stored at −20°C.

Dried peptides were resuspended in 0.1% aqueous formic acid and subjected to LC–MS/MS analysis using a Orbitrap Fusion Lumos Mass Spectrometer fitted with an EASY-nLC 1200 (Thermo Fisher Scientific) and a custom-made column heater set to 60°C. Peptides were resolved using a RP-HPLC column (75μm × 36cm) packed in-house with C18 resin (ReproSil-Pur C18–AQ, 1.9 μm resin; Dr. Maisch, r119.aq.) at a flow rate of 0.2 μLmin-1. The following gradient was used for peptide separation: from 5% B to 12% B over 5 min to 35% B over 40 min to 50% B over 15 min to 95% B over 2 min followed by 18 min at 95% B. Buffer A was 0.1% formic acid in water and buffer B was 80% acetonitrile, 0.1% formic acid in water.

The mass spectrometer was operated in DDA mode with a cycle time of 3 seconds between master scans. Each master scan was acquired in the Orbitrap at a resolution of 120,000 FWHM (at 200 m/z) and a scan range from 375 to 1500 m/z followed by MS2 scans of the most intense precursors in the linear ion trap at “Rapid” scan rate with isolation width of the quadrupole set to 1.4 m/z. Maximum ion injection time was set to 50ms (MS1) and 35 ms (MS2) with an AGC target set to 1e6 and 1e4, respectively. Only peptides with charge state 2 – 5 were included in the analysis. Monoisotopic precursor selection (MIPS) was set to Peptide, and the Intensity Threshold was set to 5e3. Peptides were fragmented by HCD (Higher-energy collisional dissociation) with collision energy set to 35%, and one microscan was acquired for each spectrum. The dynamic exclusion duration was set to 30s.

The acquired raw-files were imported into the Progenesis QI software (v2.0, Nonlinear Dynamics Limited), which was used to extract peptide precursor ion intensities across all samples applying the default parameters. The generated mgf-file was searched using MASCOT against a murine database (consisting of 33968 forward and reverse protein sequences downloaded from Uniprot on 20180710), spiked with the sequence of his-tagged *Ppargc1a* and 392 commonly observed contaminants using the following search criteria: full tryptic specificity was required (cleavage after lysine or arginine residues, unless followed by proline); 3 missed cleavages were allowed; carbamidomethylation (C) was set as fixed modification; oxidation (M) and acetyl (Protein N-term) were applied as variable modifications; mass tolerance of 10 ppm (precursor) and 0.6 Da (fragments). The database search results were filtered using the ion score to set the false discovery rate to 1% on the peptide and protein level, respectively, based on the number of reverse protein sequence hits in the dataset. Quantitative analysis results from label-free quantification were processed using the SafeQuant R package v.2.3.2^31^ (https://github.com/eahrne/SafeQuant/) to obtain peptide relative abundances. This analysis included global data normalization by equalizing the total peak/reporter areas across all LC-MS runs, data imputation using the knn algorithm, summation of peak areas per protein and LC-MS/MS run, followed by calculation of peptide abundance ratios. Only isoform specific peptide ion signals were considered for quantification. To meet additional assumptions (normality and homoscedasticity) underlying the use of linear regression models and t-Tests, MS-intensity signals were transformed from the linear to the log-scale. The summarized peptide expression values were used for statistical testing of between condition differentially abundant peptides. Here, empirical Bayes moderated t-Tests were applied, as implemented in the R/Bioconductor limma package (http://bioconductor.org/packages/release/bioc/html/limma.html), with significance defined as p-value < 0.05 and fold change ≥ 1.2. Overlap between different datasets was determined with Venny (version 2.1, https://bioinfogp.cnb.csic.es/tools/venny/). GO analysis was performed with DAVID 6.8 (https://david.ncifcrf.gov/), with significance defined as adjusted (Benjamini) p-value < 0.05. Heat maps were generated with Morpheus (https://clue.io/morpheus).

### In vitro UV crosslinking affinity purification sequencing (in vitro uvAP-seq)

Cells were transduced as for in vitro AP-MS. Total RNA was then extracted with Direct-zol(tm) RNA MiniPrep kit (Zymo Research, #R2050) and 5 μg of RNA were used for input, pull-down or negative control samples. Pull-down samples were prepared by incubating 5 μg of RNA with 12 μg of recombinant N-terminal His-Tag CTD of mouse PGC-1α (US Biological Life Sciences, #156296) for 30 min at 37°C with shaking at 500 rpm in 500 μl of PBS with RNasin® Ribonuclease Inhibitors (Promega, #N261B). Negative control samples comprised 5 μg of RNA combined with HisPur Cobalt Resin (see details below), which were incubated as described for pull-down samples. RNA-protein interactions were crosslinked by UV irradiating (5 mJ/cm^2^ at 250 nm UV wavelength) opened tubes with samples on ice, following which PBS with RNasin® Ribonuclease Inhibitors was added to 1 ml final volume. RNA-protein complexes were purified with Pierce(tm) His Protein Interaction Pull-Down Kit (Thermo Fisher Scientific, #21277) with the following modifications. HisPur Cobalt Resin was washed 5 times with wash solution and once with PBS in a 1.5 ml tube. Crosslinked samples were combined with HisPur Cobalt Resin and were incubated for 1 hour at 4°C with rotation in 1.5 ml tubes. Following incubation, HisPur Cobalt Resin containing the immobilized bait protein-RNA complexes were washed 5 times with wash buffer by centrifugation at 1250 × g for 1 min. RNA was then partially digested by re-suspending washed HisPur Cobalt Resin in 1 ml of PBS containing 10 μl of 1:1500 diluted Ambion(tm) RNase I (Thermo Fisher Scientific, #AM2294), following incubation at 37°C for 3 min with shaking at 700 rpm and then washed 5 times as described above. Subsequently, DNA was digested by incubating pull-down, negative control and input samples for 30 min at 37°C with shaking at 800 rpm with 0.05 U/µl of RQ1 RNase-Free DNase in 1X RQ1 RNase-Free DNase Reaction Buffer (Promega, # M6101). RNA was eluted from HisPur Cobalt Resin by incubating samples in a final volume of 100 µl of PBS with 0.4 mg/ml of Proteinase K (Macherey-Nagel, #740506) and incubated for 2 h at 65°C with shaking at 800 rpm. Finally, RNA was purified with Direct-zol(tm) RNA MiniPrep kit and stored at −80°C.

Purified RNA from input and pull-down samples was used to prepare libraries with Stranded Total RNA Prep with Ribo-Zero Plus kit (Illumina, # 20040525). Of note, the amount of RNA recovered from negative control samples was not sufficient for library preparation. Single-end sequencing was performed using the NextSeq 500 (Illumina) and data was analysed on the Galaxy platform (https://usegalaxy.eu/). Reads were trimmed with Trim Galore! (Galaxy version 0.4.3.1) and quality was assessed using FastQC (Galaxy version 0.72+galaxy1). Reads were aligned to the mm10 version of the mouse genome using STAR (Galaxy version 2.7.2b) and duplicated reads were removed with MarkDuplicates (Galaxy version 2.18.2.2). Next, we used MACS2 callpeak (Galaxy version, #2.1.1.20160309.6) for peak calling (q-value < 0.05 and fold change ≥ 1.5), while ChIPseeker (Galaxy version, #1.18.0+galaxy1) was used to annotate peaks. FASTA files from significant peaks were generated with Extract Genomic DNA (Galaxy version, #3.0.3) and DNA was converted to RNA with RNA/DNA (Galaxy version, # 1.0.2). FASTA files were used for motif discovery with DREME (Version 5.1.1, http://meme-suite.org/), with E-value threshold < 0.05. Normalized BAM files were generated with bamCoverage (Galaxy version 3.0.2.0) and data was visualized on the Integrated Genome Browser-9.1.4^35^ to generate representative genome browser figures. The overlap between in vitro uvAP-seq and PGC-1α ChIP-seq data (GEO accession: GSE51178) was performed with bedtools Intersect intervals (Galaxy version, #2.29.0), while overlap with RNA-seq and alternative splicing data was defined with Venny (version 2.1, https://bioinfogp.cnb.csic.es/tools/venny/).

### Confocal microscopy

Cells were seeded at 2 × 10^4^ cells per well on glass coverslips in 24 well plates and transfected as described above. Cells were fixed with 4% formaldehyde in PBS for 15 min, washed three times with PBS and incubated for 15 min with 1 µg/ml of Hoechst 33342 (Thermo Fisher Scientific, #H3570) in PBS. Next, cells were washed three times with PBS and mounted in 5 µl of ProLong(tm) Gold Antifade Mountant (Thermo Fisher Scientific, #P36930).

For immunofluorescence staining, after fixation, cells were permeabilized with 0.1% Triton X-100 in PBS for 5 min, washed three times with PBS and blocked with 10% goat serum in PBS for 30 min. Next, cells were incubated for 1 h with primary antibodies (dilution in 10% goat serum in PBS) to p-Pol 2(S5) (1:1000 dilution; Abcam, #ab5408) and H3K27ac (1:2000; Abcam, #ab4729). Following three washes with PBS, cells were incubated for 1 h with secondary antibodies (1:1000 dilution in 10% goat serum in PBS) conjugated to Alexa Fluor 568 (Thermo Fisher Scientific, #A-21124 or #A-11011). Cells were then washed three times with PBS and mounted in 5 µl of ProLong(tm) Gold Antifade Mountant with DAPI (Thermo Fisher Scientific, # P36931). All immunofluorescence staining steps were performed at room temperature.

Samples were imaged with an Olympus SpinSR CSU-W1 confocal microscope equipped with UPLAPO 100XOHR objective lens and Hamamatsu Flash4 V3 sCMOS camera. Z-stack images at 100X magnification were taken using Olympus cellSens Dimension software.

### Live-cell imaging

Cells were seeded at 2 × 10^4^ cells per chamber on an 8 well chamber slides (ibidi, #80826) and transfected as described above. Live-cells imaging was performed using FEI MORE microscope, with cells maintained at 37°C and 5% CO_2_. As light source for GFP imaging a spectraX LED with 470/24 nm excitation and 517/20 nm emission filter was used. Images were recorded with Hamamatsu OCRA flash 4.0 cooled sCMOS at 100X (numerical aperture 1.4, oil) magnification with Live Acquisition 2.5 software, while post-processing and deconvolution were carried out with Fiji 1.52p and Huygens Professional 19.10 software, respectively.

Cells were imaged before and after treatment with 5% 1,6-Hexanediol (Sigma, #240117) diluted in growth medium for up to 60 s.

Fluorescence recovery after photobleaching (FRAP) was performed with 488 nm laser and 470/24 nm filter with laser power set to 100 % (Dwell time: 0.962 ms/µm2, line overlapping 75%, ROI loop count: 10, exposure time: 20 ms, cycle time: 185 ms). Ten images were obtained pre-bleaching, while 399 images were obtained every 185 ms post-bleaching. Intensities of FRAP regions were extracted with Fiji 1.52p software tool. FRAP Data were background corrected and analysed with the online tool EasyFRAP-web (https://easyfrap.vmnet.upatras.gr/).

### Chromatin immunoprecipitation sequencing (ChIP-seq) analysis

Previously published PGC-1α ChIP-seq data (GEO accession: GSE51178)^36^ was analysed on the Galaxy platform (https://usegalaxy.eu/) for comparison with the datasets generated in this study. Reads were trimmed with Trim Galore! (Galaxy version 0.4.3.1) and quality was assessed using FastQC (Galaxy version 0.72+galaxy1). Reads were aligned to the mm10 version of the mouse genome using Bowtie2 (Galaxy Version 2.3.4.3+galaxy0) and low quality reads (phred < 20) were filtered out with Filter (Galaxy Version 2.4.1). Next, we used MACS2 callpeak (Galaxy version, #2.1.1.20160309.6) for peak calling (q-value < 0.05 and fold change ≥ 2), while ChIPseeker (Galaxy version, #1.18.0+galaxy1) was used to annotate peaks. Normalized BAM files were generated with bamCoverage (Galaxy version 3.0.2.0).

### Statistics and reproducibility

All qPCR, immunoblotting, confocal microscopy and live-imaging experiments were performed at least three independent times with similar results. FRAP quantification was performed with a total of 50 foci from independent experiments. Number of replicates per experiment is indicated in the figure legend when appropriate.

For qPCR and oxygen consumption data analysis, statistical significance was determined with one-way ANOVA with Tukey’s post hoc test using GraphPad Prism (v.8.0). Significance was considered with a p-value < 0.05. Values are expressed as mean ± SD.

RNA-seq, ChIP-seq, mass spectrometry, in vitro AP-MS and in vitro AP-seq experiments were performed once with two or three biological replicates as indicated in figure legends. Statistical analysis of these experiments is described above in their corresponding sections.

## Data availability

RNA-seq and in vitro AP-seq data will be deposited to the Gene Expression Omnibus. Whole proteome analysis and in vitro AP-MS will be deposited to PRIDE. All other data are available from the corresponding author upon reasonable request.

## Acknowledgements

We thank C. Beisel (Genomics Facility Basel, ETH Zürich), P. Demougin (Life Sciences Training Facility, Biozentrum) and K. Schleicher (Imaging Core Facility, Biozentrum) for technical help. This work was supported by the Novartis Foundation for Medical-Biological Research (J.P.S), Research Fund of the University of Basel (J.P.S), the Swiss National Science Foundation, the European Research Council (ERC) Consolidator grant 616830-MUSCLE_NET, Swiss Cancer Research grant KFS-3733-08-2015, the Swiss Society for Research on Muscle Diseases (SSEM), SystemsX.ch, the Novartis Stiftung für Medizinisch-Biologische Forschung and the University of Basel (C.H.).

## Author Contributions

JPS and CH conceived, designed and supervised the study. JPS, BK, KSH, VA, JD, GM, EVF, BKC, DR and AS performed experiments. JPS, KSH, BK, VA, TS, DR, AS, MH, SH and CH performed data analysis and interpretation. JPS and CH wrote the manuscript. All authors reviewed the manuscript.

## Competing interest

The authors declare no competing interests.

## Notes

### Competing Interest Statement

The authors have declared no competing interest.

